# Impact of sickle cell trait hemoglobin in *Plasmodium falciparum*-infected erythrocytes

**DOI:** 10.1101/2023.07.28.551025

**Authors:** Zakaria Seidu, Michael F. Ofori, Lars Hviid, Mary Lopez-Perez

**Affiliations:** Centre for Medical Parasitology, Department of Immunology and Microbiology, Faculty of Health and Medical Sciences, University of Copenhagen, Denmark; Department of Immunology, Noguchi Memorial Institute for Medical Research, College of Health Sciences, University of Ghana, Accra, Ghana; West Africa Centre for Cell Biology of Infectious Pathogens, Department of Biochemistry, Cell and Molecular Biology, University of Ghana, Accra Ghana; Centre for Medical Parasitology, Department of Infectious Diseases, Rigshospitalet, Copenhagen, Denmark

**Keywords:** HbAS, malaria, oxygen tension, PfEMP1, *pfsa*, *Plasmodium falciparum*, sickle cell trait.

## Abstract

Sickle cell trait (HbAS) confers protection against severe *Plasmodium falciparum* malaria but has little effect on infection rates *per se*. The reason for this is not fully understood. However, it appears to involve impaired parasite survival at the low oxygen tensions prevailing in the postcapillary venules where *P. falciparum*-infected erythrocytes (IEs) often accumulate. This IE sequestration is mediated by parasite-encoded IE surface ligands, primarily PfEMP1. Different variants of this family of proteins bind to host receptors with different tissue distributions. We hypothesized that *P. falciparum* parasites modulate PfEMP1 expression to enhance their survival by altering IE tissue distribution in HbAS hosts. To test this, we studied PfEMP1 expression in parasites maintained in vitro in HbAS and HbAA erythrocytes. We found that parasite survival and PfEMP1 expression were reduced in HbAS IEs, particularly at low oxygen tensions, without obvious qualitative differences in PfEMP1 expression between HbAA and HbAS IEs. In contrast, parasites growing in HbAS erythrocytes increased their transcription of *pfsa2*, a parasite gene hypothesized to be under HbS-dependent selection. Taken together, our findings support the hypothesis of quantitative but not qualitative modulation of PfEMP1 expression as a parasite strategy for coping with HbAS-related host resistance. Moreover, it provides a hint at the role of *Pfsa2* in parasite adaptation to HbAS and highlights the importance of further research.

## Introduction

The frequency of sickle hemoglobin (HbS) allele in Africa is strikingly overlapping with the present or historical distribution of *Plasmodium falciparum* (1), which is responsible for the majority of malaria cases, including severe disease with high mortality rates (2). Homozygous individuals (HbSS) develop sickle cell disease with increased morbidity and mortality, whereas heterozygous carriers (HbAS) are generally asymptomatic. HbAS also provides substantial protection against severe *P. falciparum* malaria (> 90%) and a 60% reduction in overall mortality but does not protect against *P. falciparum* infection *per se* (3). The protective mechanisms are not fully understood but appear to involve the inability of HbAS erythrocytes to sustain parasite growth at the low oxygen (O_2_) tensions prevailing in most post-capillary venules that are a preferred infected erythrocyte (IE) sequestration site (4–6). Reduced adhesion of HbAS IEs to microvascular endothelial cells has also been proposed to contribute to the protection against severe malaria (7–11).

The ability of IEs to bind to the vascular endothelium (cytoadhesion/sequestration) or to surrounding uninfected erythrocytes (rosetting) enables parasites to circumvent their destruction in the spleen. It also contributes to pathogenesis by causing circulatory obstruction and tissue inflammation. Both processes are mediated by *P. falciparum* erythrocyte membrane protein 1 (PfEMP1) family members expressed on the IE surface and encoded by ∼60 *var* genes per parasite genome (12–14). Due to allelic exclusion, only a single PfEMP1 variant is expressed per IE at a time (15). The parasites can switch transcription among the different *var* genes, thereby changing IE surface expression of PfEMP1 to evade specific immunity (16) and altering IE affinity for specific host endothelial receptors such as CD36 (17), ICAM-1 (18, 19), EPCR (19, 20), and CSA (21). IE adhesion to CD36 is generally a characteristic of uncomplicated infections (22). In contrast, adhesion to ICAM-1 and EPCR has been associated with severe malaria in children (19, 20) and adhesion to CSA with placental malaria (21). Thus, abnormal and/or reduced display of PfEMP1 on HbAS IEs (8, 10, 23) could limit their sequestration, reducing pathology and promoting parasite clearance. However, the fact that HbAS protects much less against asymptomatic and uncomplicated low-density infection than against severe disease (24) also suggests qualitative differences in PfEMP1 expression in HbAA and HbAS, respectively. On this basis, we hypothesized that *P. falciparum* parasites are able to qualitatively modulate PfEMP1 expression in HbAS to selectively avoid IE sequestration in the low O_2_ tension tissues (**Fig. S1**) that can cause severe pathology but also impairs parasite survival in HbAS IEs (4–6). To test this, we evaluated the PfEMP1 expression and the *var* gene profiles of parasites grown in vitro in HbAA and HbAS erythrocytes under several O_2_ tensions.

## Methods

### Ethical statement

The study was approved by the Ethics Review Committee (ERC) of the Ghana Health Service (GHS; GHS-ERC 008/07/19) and by the Noguchi Memorial Institute for Medical Research (NMIMR) Institutional Review Board (Federalwide Assurance FWA 00001824, NMIMR-IRB CPN 006/19). Declarations of voluntary blood sample donation and informed consent were obtained from all donors. Experiments were carried out at the University of Copenhagen (UCPH), and each participant was assigned an anonymized code.

### Blood samples collection

CPDA-anticoagulated venous blood samples were collected from healthy adult Ghanaian donors with HbAA and HbAS confirmed by Hb electrophoresis. HbAA and HbAS samples were simultaneously collected at NMIMR and immediately shipped to UCPH at 4°C. After the arrival, the cells were washed in incomplete RPMI-1640 medium to remove the buffy coat and stored at 50% hematocrit at 4°C until use (within two weeks after donation). In each experiment, HbAA and HbAS samples collected the same day were tested in parallel.

### Malaria parasite culture

Long-term in vitro-adapted *P. falciparum* clones 3D7, IT4/FCR3, and HB3, including pVBH- transfected variants (25), were maintained in culture as described previously (26, 27). In brief, parasites were grown at 5% hematocrit in serum-free RPMI-1640 medium and maintained at 37°C under controlled atmospheric conditions (2% O_2_, 5% CO_2_, 93% N_2_). The genotypic identity of the parasites (28) and the absence of *Mycoplasma* contamination using the MycoAlert Mycoplasma Detection Kit (Lonza) were verified regularly. Parasites used in the assays were maintained in culture under controlled atmospheric conditions using different O_2_ levels and 5% CO_2_ balanced with N_2_. For this, we used a C-Chamber Incubator Subchamber (BioSpherix Ltd, USA) connected to an oxygen and carbon dioxide controller (BioSpherix Ltd, C21, USA),

### Selection of IEs for surface expression of specific PfEMP1 variants

Late-stage IT4 IEs were selected for surface expression of PfEMP1 variants IT4VAR04, IT4VAR09, or IT4VAR60 using protein A-coupled DynaBeads coated with the human monoclonal antibody PAM1.4 (29, 30) or specific rabbit antisera (31) as described (32). HB3 IEs were similarly selected for the surface expression of HB3VAR06 using a specific rabbit antiserum (31). Transcription of the relevant *var* genes and IE surface expression of the corresponding PfEMP1 protein were monitored by real-time qPCR (33) and flow cytometry (32), respectively.

### Parasite growth assay

Hemozoin-containing late-stage IEs of IT4 and HB3 were purified by magnetic-activated cell sorting (MACS) as described (34). Uninfected HbAA or HbAS erythrocytes were used to adjust the parasitemia (0.5%) and hematocrit (2%) of the purified IEs, and the cultures maintained at 37°C in 2% or 19% O_2_ for six days. Spent culture medium was replaced daily. Parasitemia and parasite stage distribution (rings, trophozoites, schizonts) were evaluated daily by microscopy using Giemsa-stained thin smears. Parasitemia in HbAS was calculated relative to the corresponding HbAA (maximum parasitemia). The relative values were used to create growth curves and calculate the corresponding area under the curve (AUC).

### PfEMP1 expression on the IE surface

Parasite clones selected for expression of specific PfEMP1 as above were cultured in parallel in HbAS or HbAA erythrocytes for one cycle at 37°C under 2%, 5%, or 19% O_2_ levels and 5% CO_2._ Purified late-stage IEs were labeled with the above-mentioned antibodies, followed by FITC-conjugated anti-human IgG (Jackson ImmunoResearch) or FITC-conjugated anti-rabbit IgG (Vector Laboratories) as appropriate, and parasite nuclei were stained with Hoechst 33342 (10 µg/mL; Invitrogen). Median fluorescence intensities (MFI) of the IEs was measured using a CytoFLEX S (Beckman Coulter Life Sciences) flow cytometer and FlowLogic software (version 8.3; Inivai Technologies, Australia) (34). The average MFI in 2-3 HbAA donors per assay was considered as maximum expression (100%) and used to normalize the expression on HbAS IEs cultured in parallel.

### Rosetting assays

IEs selected as above for expression of HB3VAR06, IT4VAR09, and IT4VAR60 were maintained in HbAS or HbAA erythrocytes at 2% hematocrit in 10% human serum RPMI-1640 medium for one cycle at 5% O_2_ (5% CO_2_, 90% N_2_). Late-stage IEs were stained with ethidium bromide and the rosetting frequencies assessed by counting 200 ethidium bromide-stained IEs as described (35). Rosettes were defined as IEs having two or more adhering uninfected erythrocytes and rosetting rates in HbAS were calculated relative to the corresponding HbAA (control for maximum rosetting, 100%).

### Gene transcription analysis

Purified late-stage pVBH-transfected IEs cultured as above were collected two weeks after release from blasticidin drug pressure. The parasitemia (0.5%) and hematocrit (4%) were adjusted with uninfected HbAA or HbAS erythrocytes, and the cultures maintained at 5% O_2_ for four weeks with regular replacement of spent media and reduction of parasitemia with matching erythrocytes. Total RNA was prepared from synchronous ring-stage IEs using TRIzol (Ambion). Genomic DNA was removed by DNase I treatment (Invitrogen), and cDNA generated using SuperScript II reverse transcriptase and random primers (Invitrogen) according to the manufacturer’s instructions. QuantiTect SYBR Green PCR Master Mix (Qiagen) and three validated sets of specific primers (36–39) were used to assess *var* gene transcription profiles by real-time qPCR on a Rotorgene RG-3000 thermal cycler (Corbett Research). Additional primers were used to detect transcript levels of three non-*var* genes (*pfsa1, pfsa2, pfsa3*) (40). Individual Ct values were normalized against *seryl-tRNA synthetase* and used to determine the relative gene expression by the 2^−ΔΔCt^ method (41). Results from two independent experiments per each parasite clone are presented as the fold change relative to HbAA erythrocytes cultured in parallel.

### Statistical analysis

Data were analyzed and plotted using GraphPad Prism version 9.0 (GraphPad Software, San Diego, California, USA). The number of samples and specific statistic tests are indicated in the text or each figure. P-values < 0.05 were considered statistically significant.

## Results

### P. falciparum growth in HbAS erythrocytes is impaired at low O_2_ levels

To examine the effect of O_2_ levels in IT4 and HB3 parasite clones, we first evaluated their growth in HbAA and HbAS erythrocytes using 2% and 19% O_2_ concentrations. Over the course of the six-day culture, growth impairment in HbAS erythrocytes was apparent at both O_2_ tensions but less at 19% than at 2% O_2_ as demonstrated by a significantly higher area under the curve (AUC; 3.3 vs 1.9; p = 0.02 t-test), regardless of the parasite clone (**Fig. 1A-B**). The daily maturation stage (rings, trophozoites, schizonts) distribution of HbAA and HbAS IEs revealed that the growth defect was due to delayed progress through the cycle in the HbAS erythrocytes (**Fig. 1C**, **Fig. S2**). For instance, after two days at 2% O_2_, most of the IT4 parasites in HbAA are schizonts (∼60%), whereas in HbAS a similar proportion is still in trophozoites. These findings correspond well with earlier studies with other parasite clones using low and high O_2_ tensions (4–6).

**Figure 1.**
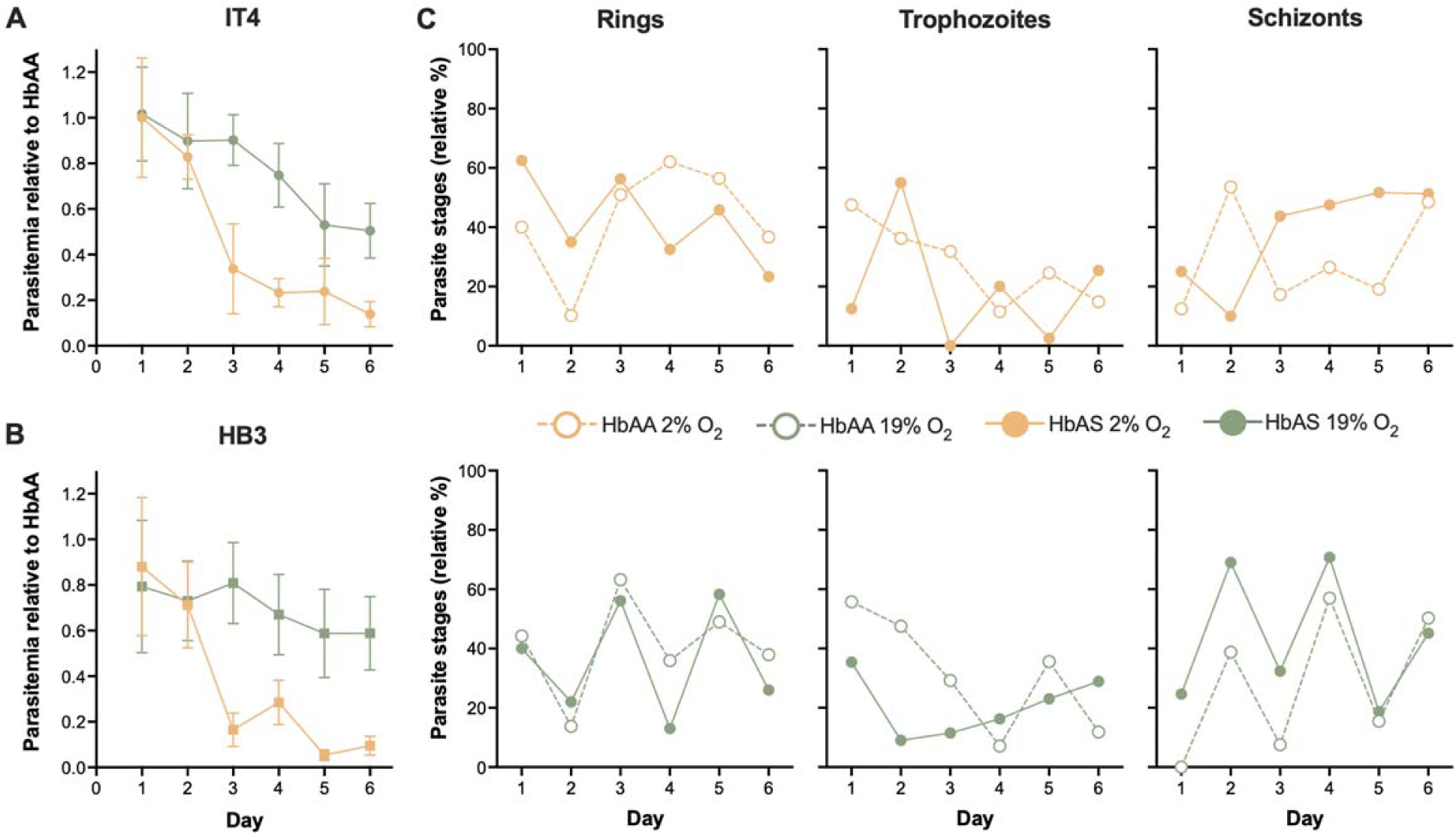
Low O_2_ levels impair intraerythrocytic parasite growth and multiplication cycle progression. Growth curves show the daily parasitemia in HbAS- relative to HbAA-IEs on the same day for IT4 (**A**) and HB3 (**B**) under 2% (orange) and 19% O_2_ levels (green). Means ± SEM of 6-7 independent assays are shown. (**C**) Relative percentage of IT4 parasite stages growing in HbAA (open symbols) and HbAS (closed symbols) erythrocytes at 2% and 19% O_2_ levels. Mean data of 6-7 independent assays are shown. Similar results for HB3 are presented in **Fig. S2**.

### PfEMP1 expression is reduced on HbAS IEs

Next, we evaluated the PfEMP1 expression of several in vitro adapted *P. falciparum* clones growing in parallel in HbAA and HbAS erythrocytes at the O_2_ tensions tested above. We also included cultures at 5% O_2_, which most closely simulates the environment in the microcirculation where IEs sequestration occurs (Supplementary Fig. 1; (42, 43)). An average of 16% reduction of PfEMP1 expression on HbAS IEs relative to HbAA IEs was observed (mean ± SEM: 84% ± 2.8%; n = 50 HbAS; p < 0.001; **Fig. 2A**), similar to what was reported previously by Cholera *et al.* (7) using ∼19% O_2_ levels (5% CO_2_ in air). Although no clear pattern relating reduction to O_2_ levels was apparent (**Fig. 2B**), the reduction varied among parasite clones selected to express specific PfEMP1 variants (**Fig. 2C-F**).

**Figure 2.**
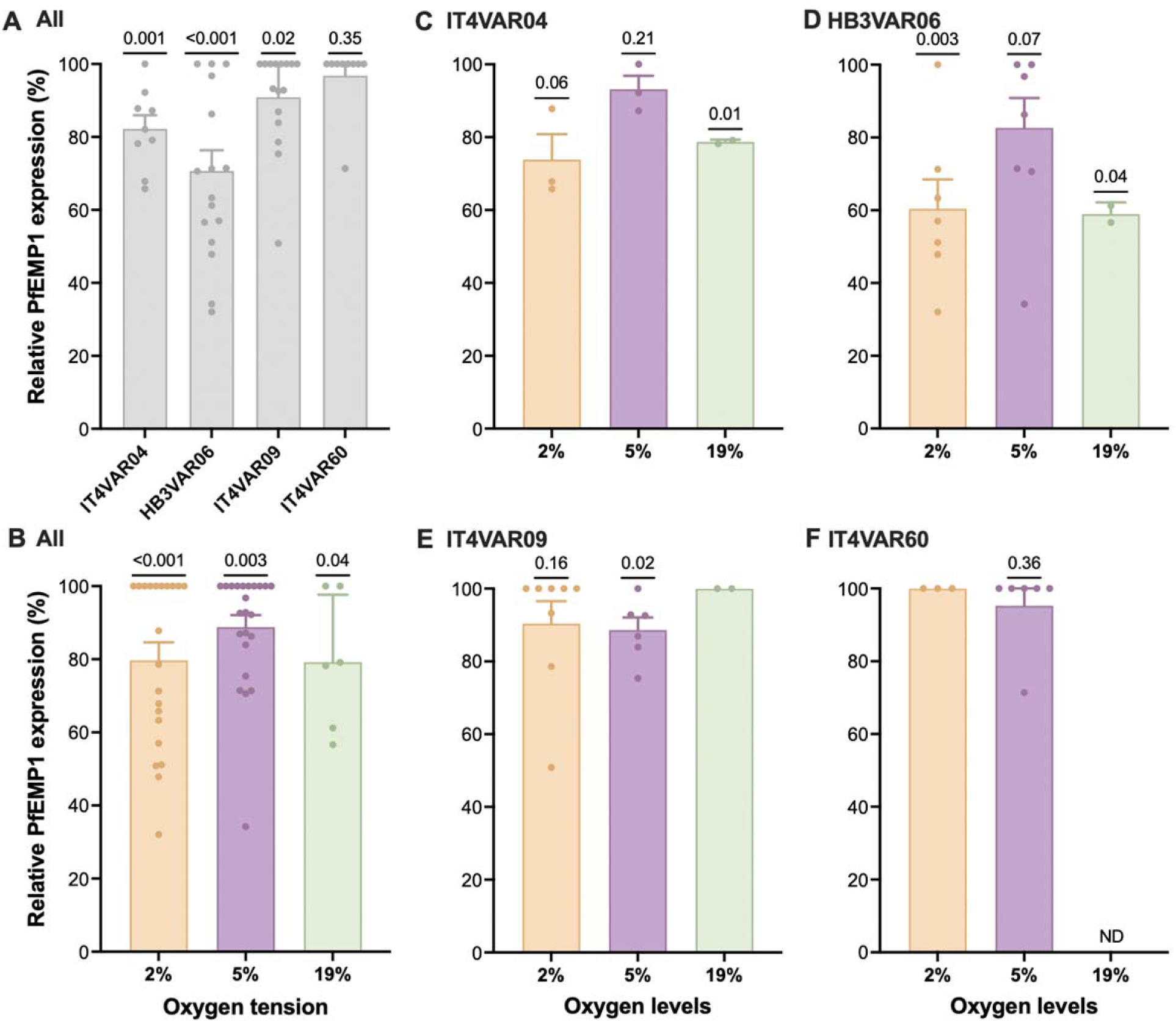
PfEMP1 expression on the surface of HbAS IEs. PfEMP1 expression on HbAS IEs relative to HbAA IEs processed in parallel at each O_2_ level, as indicated in the x-axis. Means and SEM are indicated by bars and error bars, respectively. Each symbol represents a single HbAS donor. P-values using a one-sample t-test are given. ND: no data.

The reduced PfEMP1 expression was reflected in decreased rosetting rates (76% ± 3.7; p < 0.001; n = 9 HbAS) relative to HbAA IEs for all the tested variants at 5% O_2_ (**Fig. S3**). Despite the reduced PfEMP1 protein expression, the transcription levels of the corresponding *var* genes (*hb3var06, it4var09*, and *it4var60*) in HbAS IEs were not significantly different from those in HbAA IEs at the same O_2_ levels (**Fig. S4)**.

### *var* gene transcription profiles are modified in HbAS IEs

To examine whether HbAS causes qualitative changes in the *var* gene transcription profiles, we used pVBH-transfected parasites, in which the *var* gene epigenetic memory had been erased by *bsd* transfection and drug selection. After four weeks in culture at 5% O_2_, the *var* gene transcription profiles of IT4, HB3, and 3D7 parasites grown in HbAS differed from those in HbAA erythrocytes, although no systematic differences could be discerned (**Fig. 3).** Overall, 37% of *var* genes in IT4, 68% in HB3, and 48% in 3D7 were upregulated in HbAS erythrocytes, but the transcription of only a single *var* gene (*pf13_0001*) in 3D7 was noticeably high in HbAS IEs (**Fig. 3 and Fig. S5**). In contrast, there was virtually no transcription of *it4var17, it4var22*, and *it4var40* by IT4 parasites in HbAS IEs, or of *hb3var09* and *hb3var20* by HB3 parasites in HbAS IEs.

**Figure 3.**
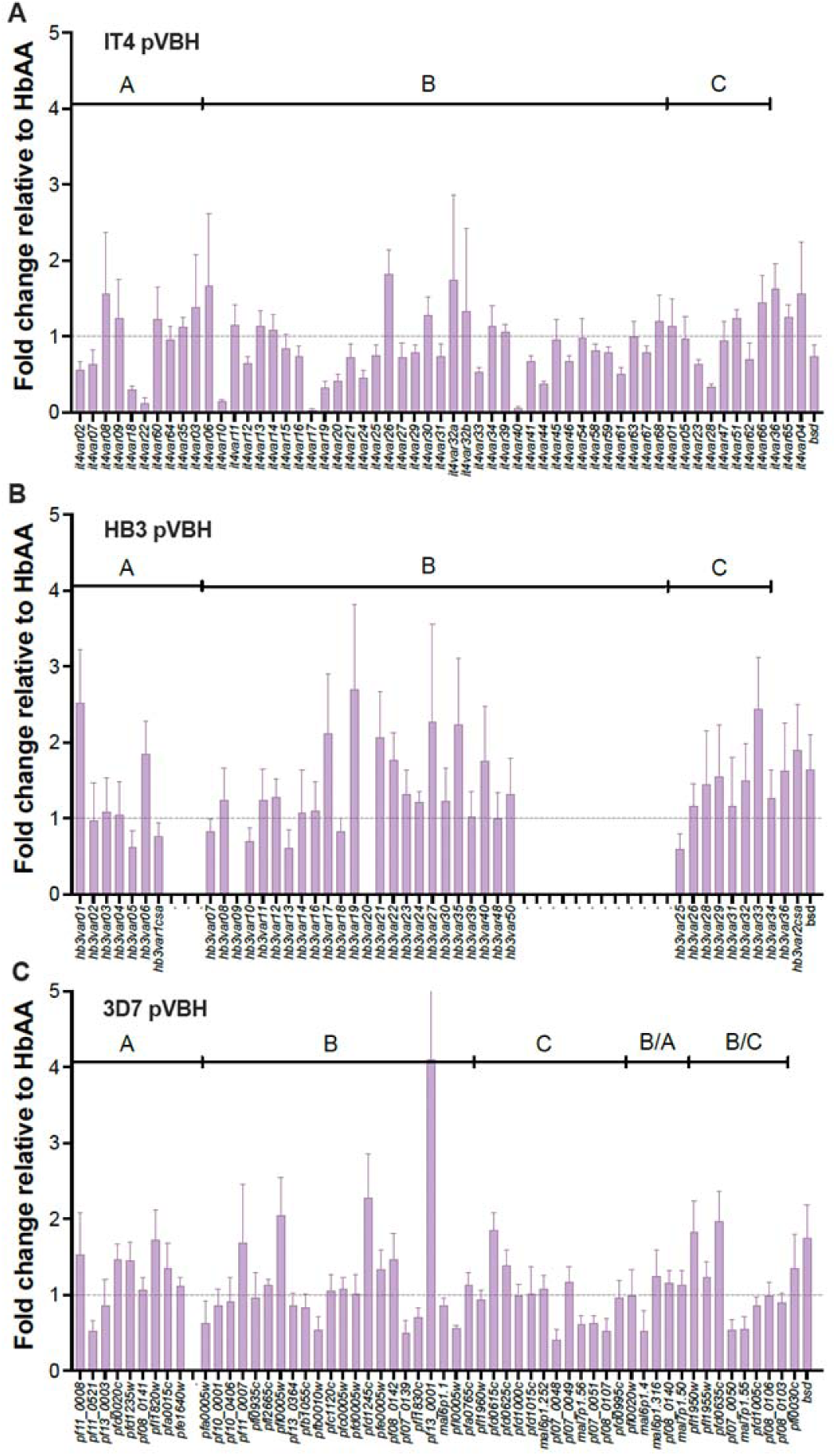
var gene transcription profiles in P. falciparum parasites growing in HbAS erythrocytes. Transcription of *var* genes in (**A**) IT4 pVBH, (**B**) HB3 pVBH, and (**C**) 3D7 pVBH growing in HbAS erythrocytes after four weeks at 5% O_2_. Fold changes relative to transcription in HbAA are shown. *var* genes are ordered according to gene group. Data are presented as means ± SEM of 5-6 HbAS individual donors.

### Pfsa2 is upregulated on HbAS IEs

It was recently reported that parasite carriage of specific *Pfsa* (*P. falciparum* sickle-associated) alleles might offset the protective effect of HbS against severe malaria (40). We therefore tested whether the transcription of *Pfsa1* (PF3D7_0215300; acyl-CoA synthetase family member PfACS8), *Pfsa2* (PF3D7_0220300; an exported protein), and *Pfsa3* (PF3D7_1127000; a putative tyrosine phosphatase) differed between HbAA and HbAS IEs. After four weeks in culture, the three genes were transcribed by IT4, HB3, and 3D7 parasites growing in both types of erythrocytes, but *Pfsa2* was upregulated in parasites growing in HbAS IEs compared to HbAA (**Fig. 4**). In contrast with the overexpression, none of the three tested parasites clones carry the mutation in the corresponding alleles as observed in the multiple sequence alignment (**Supplementary Fig S6**).

**Figure 4.**
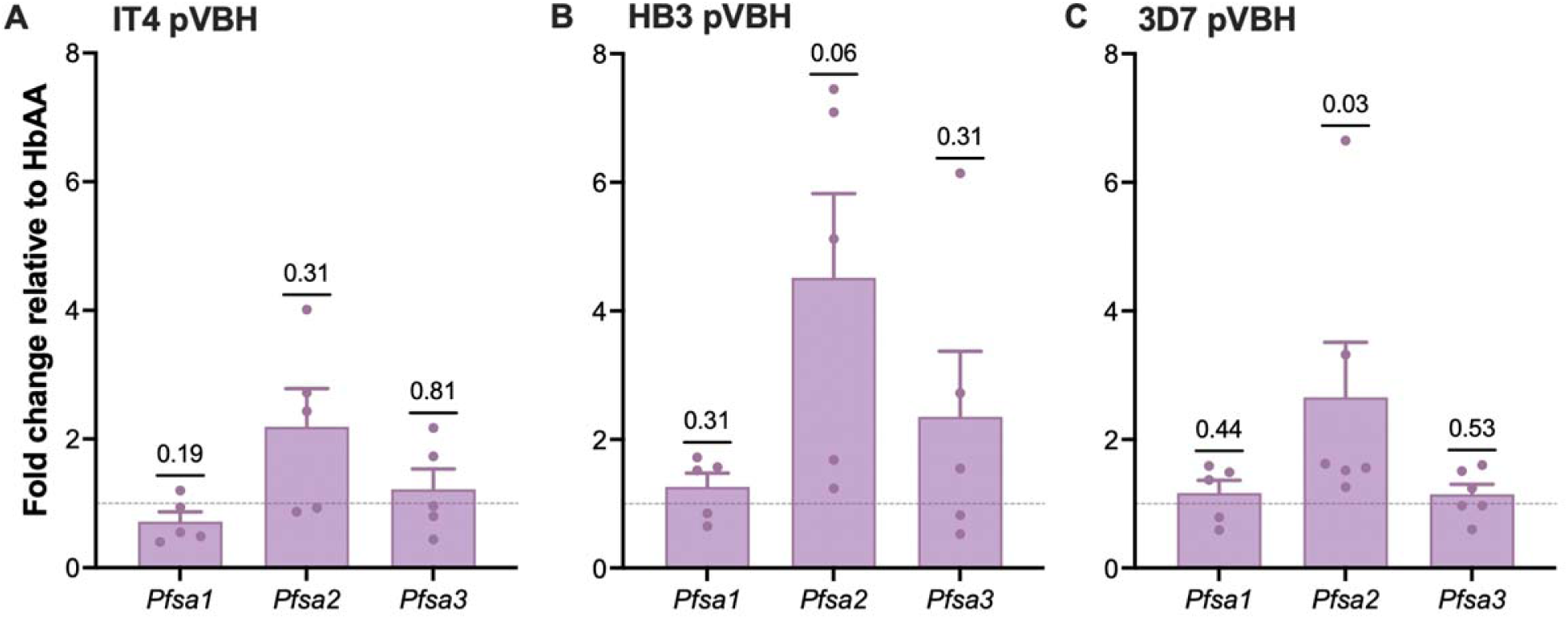
Pfsa gene transcription in P. falciparum parasites. Transcription of *Pfsa* genes in (**A**) IT4 pVBH, (**B**) HB3 pVBH, and (**C**) 3D7 pVBH growing in HbAS erythrocytes after four weeks at 5% O_2_. Fold changes relative to transcription in HbAA erythrocytes processed in parallel and p-values using Wilcoxon signed-rank test are shown. Data are presented as means ± SEM of 5-6 HbAS individual donors.

## Discussion

Individuals with sickle cell trait (HbAS) are protected from severe *P. falciparum* malaria (3). The specific mechanisms mediating this protection are less clear but likely involve restricted intraerythrocytic growth at low O_2_ tensions (4–6) and reduced adhesion of IEs to host cell receptors (7–11). Conceivably, this is sufficient to reduce the parasite burden and associated inflammation in HbAS individuals to relatively benign levels that prevent severe pathology. At the same time, it might be conducive to the persistence of low-level, relatively asymptomatic parasitemia (44), promoting the acquisition of protective immunity in the absence of evident disease (45) (*Oleinikov et al. unpublished data*).

HbAS erythrocytes collapse (“sickle”) due to the polymerization of HbS when the O_2_ tension is low, rendering them unsuitable to support *P. falciparum* growth (46). However, the possibility of parasites responding to this by qualitatively modulating PfEMP1 expression to avoid IE sequestration in tissues with low O_2_ tension has not been investigated. There is some indirect support for this idea from studies showing that HbAS does not protect against placental malaria (47, 48), characterized by selective IE sequestration in a high O_2_ tension tissue (**Fig. S1**), and here we set out to provide additional evidence.

Using three *P. falciparum* clones with divergent *var* gene repertoires (49), we could confirm previous observations (4–6) that parasite growth in HbAS IEs is impaired, particularly at low O_2_ tension. We also observed a reduced expression of PfEMP1 on the surface of IEs, particularly at low O_2_ tensions. This observation is consistent with a previous report conducted with unselected IEs (7) and a study directly comparing the number of PfEMP1 molecules on HbAA and HbAS IEs selected to express a particular variant (VAR2CSA) (10). The formation of the electron-dense surface knob on which PfEMP1 is displayed is compromised in HbAS (23), and although Sanchez *et al.* (10) found more VAR2CSA molecules per knob on HbAS IEs, the increase was more than offset by the lower number of knobs per IE. Despite the reduced expression of IE surface expression of PfEMP1, we did not find evidence of an underlying defect in the transcription of the *var* genes encoding these adhesive proteins, suggesting that the cause of the reduced PfEMP1 expression is post-transcriptional, in agreement with the conclusions of earlier work on this topic (8, 23, 50).

Overall, our data support the generally held opinion that the protective effect of HbAS is primarily the direct consequence of impaired intracellular transport and IE surface display of PfEMP1, leading to reduced cytoadhesion (7, 9) and improved splenic removal of IEs (51). Our finding of reduced rosetting rates in HbAS IEs supports this notion. After long-term culture in HbAS erythrocytes, we observed only minor changes in *var* gene transcription relative to parasites grown in HbAA erythrocytes, without any obvious consistency among the clones tested. The uncomplicated malaria-associated Group B variant *pf13_0001* predicted to bind to CD36 (DBLα0.11- CIDRα2.4) was highly transcribed among parasites in HbAS IEs, whereas transcription of the severe malaria-associated variants *it4var22* (20) and *hb3var09* (33) was virtually absent. Those results agree with our observation that children with HbAS have a lower IgG-specific response than those with HbAA to PfEMP1 variants belonging to groups A and B/A (*Oleinikov et al. under review*). Nevertheless, these observations need further validation to support the alternative or complementary hypothesis that *P. falciparum* parasites in HbAS IEs can modulate which PfEMP1 variants are expressed to avoid IE sequestration in tissues with low O_2_ tension.

Our observations partly agree with a recent study, which did not observe marked changes in *var* gene transcription of 3D7 parasites grown in HbAS erythrocytes for a single multiplication cycle (11). In a separate study, the same team identified hundreds of non-*var* transcripts that differed between parasites grown in HbAS and HbAA erythrocytes (50). These were mainly related to folding machinery, oxidative stress response, and protein export machinery. As HbAS does not affect the redox potential of IEs (52), oxidative stress may not be important for its malaria-protective function. However, changes were also found among genes encoding proteins in Maurer’s clefts, parasite-derived membranous structures associated with PfEMP1 trafficking and described as amorphous and less functional in HbAS IEs (53). A recent genome-wide association study reported that protection against severe malaria in children with HbS also depends on parasite genotype, involving variants at three loci named *Pfsa (Pfsa1* chr2:631,190 T > A*, Pfsa2* chr2:814,288 C > T, and *Pfsa3* chr11:1,058,035 T > A and chr11:1,057,437 T > C) (40). The protein encoded by *Pfsa2* appears integrated into the Maurer’s clefts and is perhaps associated with the PfEMP1 expression on the IEs surface (54), whereas the *Pfsa3* product has been observed in the food vacuole (55). Band et al. (40) reported that *Pfsa3* gene was overexpressed in children with HbAS compared with those with HbAA and found that the specific single nucleotide polymorphisms probably explain this increased expression (40). We observed that *Pfsa1, Pfsa2, and Pfsa3* were all transcribed by parasites in either erythrocyte type, in accordance with the Band *et al.* study (40). Despite the parasites tested in this study being wildtype for the corresponding allele, we found that *Pfsa2* transcription was upregulated in HbAS IEs after four weeks in culture, suggesting a potential involvement in a compensatory mechanism for the dysfunctional PfEMP1 trafficking described in HbAS IEs (23, 53). This deserves further study beyond the scope of the current work.

Our in vitro study clearly has limitations and may not wholly reflect the in vivo conditions in which sickling, acquired PfEMP1-specific immunity (*Oleinikov et al. under review*), and the adequate splenic clearance of HbAS IEs (51) are likely to contribute to controlling parasite growth in individuals with HbAS. Furthermore, our study was focused on *var* gene transcription and its relation to O_2_ tension; thus, the involvement of additional genes (50) under other conditions cannot be ruled out. Finally, the impact of mutations in the transcriptional changes in *Pfsa2* was not analyzed.

Using several *P. falciparum* clones, our study confirms the negative effect of low O_2_ tension on intraerythrocytic parasite growth and the impact of HbAS on PfEMP1 IE surface expression. Whereas the results with *Pfsa* gene are inconclusive, they hint at the role of *Pfsa2* in parasite adaptation to HbAS and highlight the importance of further research. Altogether, our study adds to the evidence that the protective effect of HbAS against severe *P. falciparum* malaria is multifactorial.

## Acknowledgments

We thank all the Ghanaian blood donors. We also thank Ron Dzikowski (The Hebrew University of Jerusalem, Israel) and Kirk Deitsch (Weill Medical College of Cornell University, NY) for providing the pVBH-transfected 3D7 clone and the pVBH plasmid, respectively. Eric Kyei-Baffour (NMIMR, University of Ghana), Morten Pontoppidan, and Maiken Visti (University of Copenhagen) are thanked for their technical assistance.

This work was funded by the Independent Research Fund Denmark (grant 0134-00123B; LH and MLP) and Danish International Development Agency, Danida (grant 17 02 KU; LH, MO). ZS was supported by a PhD scholarship from the Danida-sponsored Building Stronger Universities initiative grant (BSUIII-UG). The funders had no role in study design, data collection and analysis, decision to publish, or preparation of the manuscript.

## Conflict of interest

The authors declare no conflict of interest.

## Author contributions

LH and MLP conceptualization. ZS and MLP methodology and investigation. MLP data curation and formal analysis. ZS and MFO resources. ZS and MLP Writing - original draft. ZS, MFO, LH and MLP Writing - review & editing. All authors approved the final version of the manuscript.

## References

1. Piel FB, Patil AP, Howes RE, Nyangiri OA, Gething PW, Williams TN, Weatherall DJ, Hay SI. 2010. Global distribution of the sickle cell gene and geographical confirmation of the malaria hypothesis. Nat Commun 1:104.

2. WHO. 2022. World Malaria Report 2022. World Health Organization, Geneva.

3. Taylor SM, Parobek CM, Fairhurst RM. 2012. Haemoglobinopathies and the clinical epidemiology of malaria: a systematic review and meta-analysis. Lancet Infect Dis 12:457–468.

4. Friedman MJ. 1978. Erythrocytic mechanism of sickle cell resistance to malaria. Proc Natl Acad Sci U S A 75:1994–1997.

5. Pasvol G, Weatherall DJ, Wilson RJ. 1978. Cellular mechanism for the protective effect of haemoglobin S against *P. falciparum* malaria. Nature 274:701–703.

6. Archer NM, Petersen N, Clark MA, Buckee CO, Childs LM, Duraisingh MT. 2018. Resistance to *Plasmodium falciparum* in sickle cell trait erythrocytes is driven by oxygen-dependent growth inhibition. Proc Natl Acad Sci U S A 115:7350–7355.

7. Cholera R, Brittain NJ, Gillrie MR, Lopera-Mesa TM, Diakite SA, Arie T, Krause MA, Guindo A, Tubman A, Fujioka H, Diallo DA, Doumbo OK, Ho M, Wellems TE, Fairhurst RM. 2008. Impaired cytoadherence of *Plasmodium falciparum*-infected erythrocytes containing sickle hemoglobin. Proc Natl Acad Sci U S A 105:991–996.

8. Kilian N, Srismith S, Dittmer M, Ouermi D, Bisseye C, Simpore J, Cyrklaff M, Sanchez CP, Lanzer M. 2015. Hemoglobin S and C affect protein export in *Plasmodium falciparum*-infected erythrocytes. Biol Open 4:400–410.

9. Lansche C, Dasanna AK, Quadt K, Frohlich B, Missirlis D, Tetard M, Gamain B, Buchholz B, Sanchez CP, Tanaka M, Schwarz US, Lanzer M. 2018. The sickle cell trait affects contact dynamics and endothelial cell activation in *Plasmodium falciparum*- infected erythrocytes. Commun Biol 1:211.

10. Sanchez CP, Karathanasis C, Sanchez R, Cyrklaff M, Jager J, Buchholz B, Schwarz US, Heilemann M, Lanzer M. 2019. Single-molecule imaging and quantification of the immune-variant adhesin VAR2CSA on knobs of *Plasmodium falciparum*-infected erythrocytes. Commun Biol 2:172.

11. Petersen JEV, Saelens JW, Freedman E, Turner L, Lavstsen T, Fairhurst RM, Diakite M, Taylor SM. 2021. Sickle-trait hemoglobin reduces adhesion to both CD36 and EPCR by *Plasmodium falciparum*-infected erythrocytes. PLoS Pathog 17:e1009659.

12. Su XZ, Heatwole VM, Wertheimer SP, Guinet F, Herrfeldt JA, Peterson DS, Ravetch JA, Wellems TE. 1995. The large diverse gene family *var* encodes proteins involved in cytoadherence and antigenic variation of *Plasmodium falciparum*-infected erythrocytes. Cell 82:89–100.

13. Rask TS, Hansen DA, Theander TG, Gorm Pedersen A, Lavstsen T. 2010. *Plasmodium falciparum* erythrocyte membrane protein 1 diversity in seven genomes--divide and conquer. PLoS Comput Biol 6.

14. Hviid L, Jensen AT. 2015. PfEMP1 - A parasite protein family of key importance in *Plasmodium falciparum m*alaria immunity and pathogenesis. Adv Parasitol 88:51–84.

15. Deitsch KW, Calderwood MS, Wellems TE. 2001. Malaria. Cooperative silencing elements in var genes. Nature 412:875–876.

16. Roberts DJ, Craig AG, Berendt AR, Pinches R, Nash G, Marsh K, Newbold CI. 1992. Rapid switching to multiple antigenic and adhesive phenotypes in malaria. Nature 357:689–692.

17. Oquendo P, Hundt E, Lawler J, Seed B. 1989. CD36 directly mediates cytoadherence of *Plasmodium falciparum* parasitized erythrocytes. Cell 58:95–101.

18. Berendt AR, Simmons DL, Tansey J, Newbold CI, Marsh K. 1989. Intercellular adhesion molecule-1 is an endothelial cell adhesion receptor for *Plasmodium falciparum*. Nature 341:57–59.

19. Lennartz F, Adams Y, Bengtsson A, Olsen RW, Turner L, Ndam NT, Ecklu-Mensah G, Moussiliou A, Ofori MF, Gamain B, Lusingu JP, Petersen JE, Wang CW, Nunes-Silva S, Jespersen JS, Lau CK, Theander TG, Lavstsen T, Hviid L, Higgins MK, Jensen AT. 2017. Structure-guided identification of a family of dual receptor-binding PfEMP1 that is associated with cerebral malaria. Cell Host Microbe 21:403–414.

20. Turner L, Lavstsen T, Berger SS, Wang CW, Petersen JE, Avril M, Brazier AJ, Freeth J, Jespersen JS, Nielsen MA, Magistrado P, Lusingu J, Smith JD, Higgins MK, Theander TG. 2013. Severe malaria is associated with parasite binding to endothelial protein C receptor. Nature 498:502–505.

21. Salanti A, Dahlback M, Turner L, Nielsen MA, Barfod L, Magistrado P, Jensen AT, Lavstsen T, Ofori MF, Marsh K, Hviid L, Theander TG. 2004. Evidence for the involvement of VAR2CSA in pregnancy-associated malaria. J Exp Med 200:1197–1203.

22. Cabrera A, Neculai D, Kain KC. 2014. CD36 and malaria: friends or foes? A decade of data provides some answers. Trends Parasitol 30:436–444.

23. Cyrklaff M, Srismith S, Nyboer B, Burda K, Hoffmann A, Lasitschka F, Adjalley S, Bisseye C, Simpore J, Mueller AK, Sanchez CP, Frischknecht F, Lanzer M. 2016. Oxidative insult can induce malaria-protective trait of sickle and fetal erythrocytes. Nat Commun 7:13401.

24. Lopera-Mesa TM, Doumbia S, Konate D, Anderson JM, Doumbouya M, Keita AS, Diakite SA, Traore K, Krause MA, Diouf A, Moretz SE, Tullo GS, Miura K, Gu W, Fay MP, Taylor SM, Long CA, Diakite M, Fairhurst RM. 2015. Effect of red blood cell variants on childhood malaria in Mali: a prospective cohort study. Lancet Haematol 2:e140–149.

25. Fastman Y, Noble R, Recker M, Dzikowski R. 2012. Erasing the epigenetic memory and beginning to switch--the onset of antigenic switching of var genes in *Plasmodium falciparum*. PLoS One 7:e34168.

26. Cranmer SL, Magowan C, Liang J, Coppel RL, Cooke BM. 1997. An alternative to serum for cultivation of *Plasmodium falciparum in vitro*. Trans R Soc Trop Med Hyg 91:363–365.

27. Lopez-Perez M, Seidu Z. 2022. Establishing and maintaining in vitro cultures of asexual blood stages of *Plasmodium falciparum*, p 37-49. *In* Jensen AR, Hviid L (ed), Malaria Immunology, Methods in Molecular Biology, vol 2470. Humana, New York, NY.

28. Snounou G, Zhu X, Siripoon N, Jarra W, Thaithong S, Brown KN, Viriyakosol S. 1999. Biased distribution of *msp1* and *msp2* allelic variants in *Plasmodium falciparum* populations in Thailand. Trans R Soc Trop Med Hyg 93:369–374.

29. Barfod L, Bernasconi NL, Dahlback M, Jarrossay D, Andersen PH, Salanti A, Ofori MF, Turner L, Resende M, Nielsen MA, Theander TG, Sallusto F, Lanzavecchia A, Hviid L. 2007. Human pregnancy-associated malaria-specific B cells target polymorphic, conformational epitopes in VAR2CSA. Mol Microbiol 63:335–347.

30. Raghavan SSR, Dagil R, Lopez-Perez M, Conrad J, Bassi MR, Quintana MDP, Choudhary S, Gustavsson T, Wang Y, Gourdon P, Ofori MF, Christensen SB, Minja DTR, Schmiegelow C, Nielsen MA, Barfod L, Hviid L, Salanti A, Lavstsen T, Wang K. 2022. Cryo-EM reveals the conformational epitope of human monoclonal antibody PAM1.4 broadly reacting with polymorphic malarial protein VAR2CSA. PLoS Pathog 18:e1010924.

31. Stevenson L, Laursen E, Cowan GJ, Bandoh B, Barfod L, Cavanagh DR, Andersen GR, Hviid L. 2015. alpha2-macroglobulin can crosslink multiple *Plasmodium falciparum* Erythrocyte Membrane Protein 1 (PfEMP1) molecules and may facilitate adhesion of parasitized erythrocytes. PLoS Pathog 11:e1005022.

32. Lopez-Perez M, Olsen RW. 2022. Immunomagnetic selection of *Plasmodium falciparum*-infected erythrocytes expressing particular PfEMP1 variants. Methods Mol Biol 2470:69–78.

33. Quintana MDP, Ecklu-Mensah G, Tcherniuk SO, Ditlev SB, Oleinikov AV, Hviid L, Lopez-Perez M. 2019. Comprehensive analysis of Fc-mediated IgM binding to the *Plasmodium falciparum* erythrocyte membrane protein 1 family in three parasite clones. Sci Rep 9:6050.

34. Lopez-Perez M, Olsen RW. 2022. Immunomagnetic selection of *Plasmodium falciparum*-infected erythrocytes expressing particular PfEMP1 variants, p 69–78. *In* Jensen AR, Hviid L (ed), Malaria Immunology, Methods in Molecular Biology, vol 2470.

35. Lopez-Perez M, van der Puije W, Castberg FC, Ofori MF, Hviid L. 2020. Binding of human serum proteins to *Plasmodium falciparum*-infected erythrocytes and its association with malaria clinical presentation. Malar J 19:362.

36. Dahlback M, Lavstsen T, Salanti A, Hviid L, Arnot DE, Theander TG, Nielsen MA. 2007. Changes in var gene mRNA levels during erythrocytic development in two phenotypically distinct *Plasmodium falciparum* parasites. Malar J 6:78.

37. Joergensen L, Bengtsson DC, Bengtsson A, Ronander E, Berger SS, Turner L, Dalgaard MB, Cham GK, Victor ME, Lavstsen T, Theander TG, Arnot DE, Jensen AT. 2010. Surface co-expression of two different PfEMP1 antigens on single *Plasmodium falciparum*-infected erythrocytes facilitates binding to ICAM1 and PECAM1. PLoS Pathog 6:e1001083.

38. Soerli J, Barfod L, Lavstsen T, Bernasconi NL, Lanzavecchia A, Hviid L. 2009. Human monoclonal IgG selection of *Plasmodium falciparum* for the expression of placental malaria-specific variant surface antigens. Parasite Immunol 31:341–346.

39. Wang CW, Lavstsen T, Bengtsson D, Magistrado P, Berger SS, Marquard MA, Alifrangis M, Lusingu J, Theander T, Turner L. 2012. Evidence for in vitro and in vivo expression of the conserved VAR3 (type 3) *Plasmodium falciparum* erythrocyte membrane protein 1. Malar J 11:129.

40. Band G, Leffler EM, Jallow M, Sisay-Joof F, Ndila CM, Macharia AW, Hubbart C, Jeffreys AE, Rowlands K, Nguyen T, Goncalves S, Ariani CV, Stalker J, Pearson RD, Amato R, Drury E, Sirugo G, d’Alessandro U, Bojang KA, Marsh K, Peshu N, Saelens JW, Diakite M, Taylor SM, Conway DJ, Williams TN, Rockett KA, Kwiatkowski DP. 2022. Malaria protection due to sickle haemoglobin depends on parasite genotype. Nature 602:106–111.

41. Livak KJ, Schmittgen TD. 2001. Analysis of relative gene expression data using real-time quantitative PCR and the 2(-Delta Delta C(T)) Method. Methods 25:402–408.

42. Carreau A, El Hafny-Rahbi B, Matejuk A, Grillon C, Kieda C. 2011. Why is the partial oxygen pressure of human tissues a crucial parameter? Small molecules and hypoxia. J Cell Mol Med 15:1239–1253.

43. Wenger RH, Kurtcuoglu V, Scholz CC, Marti HH, Hoogewijs D. 2015. Frequently asked questions in hypoxia research. Hypoxia (Auckl) 3:35–43.

44. Gong L, Maiteki-Sebuguzi C, Rosenthal PJ, Hubbard AE, Drakeley CJ, Dorsey G, Greenhouse B. 2012. Evidence for both innate and acquired mechanisms of protection from *Plasmodium falciparum* in children with sickle cell trait. Blood 119:3808–3814.

45. Williams TN, Mwangi TW, Roberts DJ, Alexander ND, Weatherall DJ, Wambua S, Kortok M, Snow RW, Marsh K. 2005. An immune basis for malaria protection by the sickle cell trait. PLoS Med 2:e128.

46. Luzzatto L, Nwachuku-Jarrett ES, Reddy S. 1970. Increased sickling of parasitised erythrocytes as mechanism of resistance against malaria in the sickle-cell trait. Lancet 295:319–321.

47. Chauvet M, Tetard M, Cottrell G, Aussenac F, Brossier E, Denoyel L, Hanny M, Lohezic M, Milet J, Ndam NT, Pineau D, Roman J, Luty AJF, Gamain B, Migot-Nabias F, Merckx A. 2019. Impact of hemoglobin S trait on cell surface antibody recognition of *Plasmodium falciparum*-infected erythrocytes in pregnancy-associated malaria. Open Forum Infect Dis 6:ofz156.

48. Lopez-Perez M, Viwami F, Seidu Z, Jensen ATR, Doritchamou J, Ndam NT, Hviid L. 2021. PfEMP1-specific immunoglobulin G reactivity among beninese pregnant women with sickle cell trait. Open Forum Infect Dis 8:ofab527.

49. Kraemer SM, Kyes SA, Aggarwal G, Springer AL, Nelson SO, Christodoulou Z, Smith LM, Wang W, Levin E, Newbold CI, Myler PJ, Smith JD. 2007. Patterns of gene recombination shape var gene repertoires in *Plasmodium falciparum*: comparisons of geographically diverse isolates. BMC Genomics 8:45.

50. Saelens JW, Petersen JEV, Freedman E, Moseley RC, Konate D, Diakite SAS, Traore K, Vance N, Fairhurst RM, Diakite M, Haase SB, Taylor SM. 2021. Impact of sickle cell trait hemoglobin on the intraerythrocytic transcriptional program of *Plasmodium falciparum*. mSphere 6:e0075521.

51. Diakite SA, Ndour PA, Brousse V, Gay F, Roussel C, Biligui S, Dussiot M, Prendki V, Lopera-Mesa TM, Traore K, Konate D, Doumbia S, Cros J, Dokmak S, Fairhurst RM, Diakite M, Buffet PA. 2016. Stage-dependent fate of *Plasmodium falciparum*-infected red blood cells in the spleen and sickle-cell trait-related protection against malaria. Malar J 15:482.

52. Haag M, Kehrer J, Sanchez CP, Deponte M, Lanzer M. 2022. Physiological jump in erythrocyte redox potential during *Plasmodium falciparum* development occurs independent of the sickle cell trait. Redox Biol 58:102536.

53. Kilian N, Dittmer M, Cyrklaff M, Ouermi D, Bisseye C, Simpore J, Frischknecht F, Sanchez CP, Lanzer M. 2013. Haemoglobin S and C affect the motion of Maurer’s clefts in *Plasmodium falciparum*-infected erythrocytes. Cell Microbiol 15:1111–1126.

54. Roling L. 2022. Investigating the importance of exported proteins for survival and host cell modification in asexual blood stages of *Plasmodium falciparum*. PhD. Ruprecht Karl University of Heidelberg, Heidelberg.

55. Lamarque M, Tastet C, Poncet J, Demettre E, Jouin P, Vial H, Dubremetz JF. 2008. Food vacuole proteome of the malarial parasite *Plasmodium falciparum*. Proteomics Clin Appl 2:1361–1374.

